# Phylogenetic analysis of TALE superclass homeobox genes in amphipod crustaceans

**DOI:** 10.1101/2021.08.10.455793

**Authors:** Wai Hoong Chang, Alvina G. Lai

## Abstract

TALE class genes are a group of developmentally conserved transcription factors found in animals. Here, we describe the identification and phylogenetic analysis of TALE class genes in amphipod crustaceans. We identified 241 putative TALE class genes from 56 amphipod crustacean species. Phylogenetic analysis of the genes revealed six subclasses. We provide a list of FASTA sequences of the genes identified. Results from this work may inform future evolutionary and comparative genomics studies on animal development.

## Introduction

Animals exhibit remarkably phenotypic diversity underpinned by genetic innovations that influence developmental processes. The specification of body plans in animals is regulated by a conserved bilaterian genetic toolkit that encodes a collection of transcription factors and their target cis-regulatory modules found within large gene regulatory networks evolving at different rates (1). These genetic precursors have been discovered in ancient clades such as in choanoflagellates and non-bilaterian animals, suggesting their role during the early evolution of bilaterians (2).

Among the developmental gene families are a highly conserved group of homeodomain-containing transcription factors known for their roles in body plan patterning. Homeodomain proteins are characterised by the presence of a conserved DNA-binding region known as the homeodomain that are seen in Homeobox proteins (3). Homeobox genes consist of a large family of genes subdivided into 11 gene classes in animals (4). Homeobox genes have been widely characterised in a range of model organisms (5). However, systematic characterisation of this gene family is currently lacking in non-model animals. Here, we present the identification and phylogenetic analyses of the three amino acid loop extension (TALE) homeobox genes in amphipod crustaceans.

## Methods

To identify TALE class genes, we previously performed BLAST searches using transcriptome datasets of 56 amphipod crustacean species retrieved from the European Nucleotide Archive (6, 7) (Figure 1). Reference sequences of TALE class genes were downloaded from the NCBI database. We performed tBLASTn searches against the crustacean transcriptomes using the reference sequences with blocks substitution matrices BLOSUM45 and BLOSUM62. BLAST results were filtered for unique hits and by e-value of < 10-5. Reciprocal BLAST was performed against the GenBank non-redundant (nr) database. Redundant transcripts having at least 98% identity were collapsed using the CD-HIT tool. The HMMER hmmscan tool was used to search against profile hidden Markov model (HMM) libraries to identify Pfam homeodomains (8). Multiple sequence alignment was performed using the MAFFT tool (9). Phylogenetic tree was constructed from the alignment using RAxML WAG + G model to generate a maximum likelihood tree (10).

**Fig. 1.**
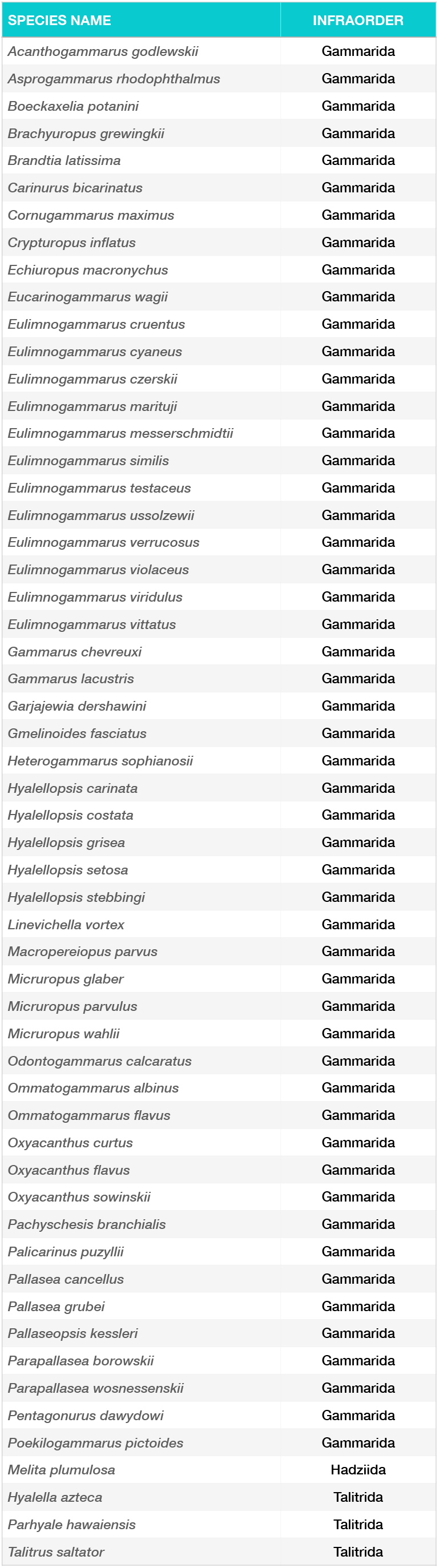
List of amphipod species analysed in this study.

## Results

We employed transcriptome datasets from 56 amphipod crustacean species. From these datasets, we identified 241 putative TALE class genes using BLAST and profile HMM homology searches. Phylogenetic analyses of TALE class genes revealed that the genes can be further divided into six subclasses: Tgif, Meis, Pknox, Irx, Mkx and Pbx. The tree topology of amphipod TALE class orthologs is depicted in Figure 2. Fasta sequences are provided as Supplementary data 1.

**Fig. 2.**
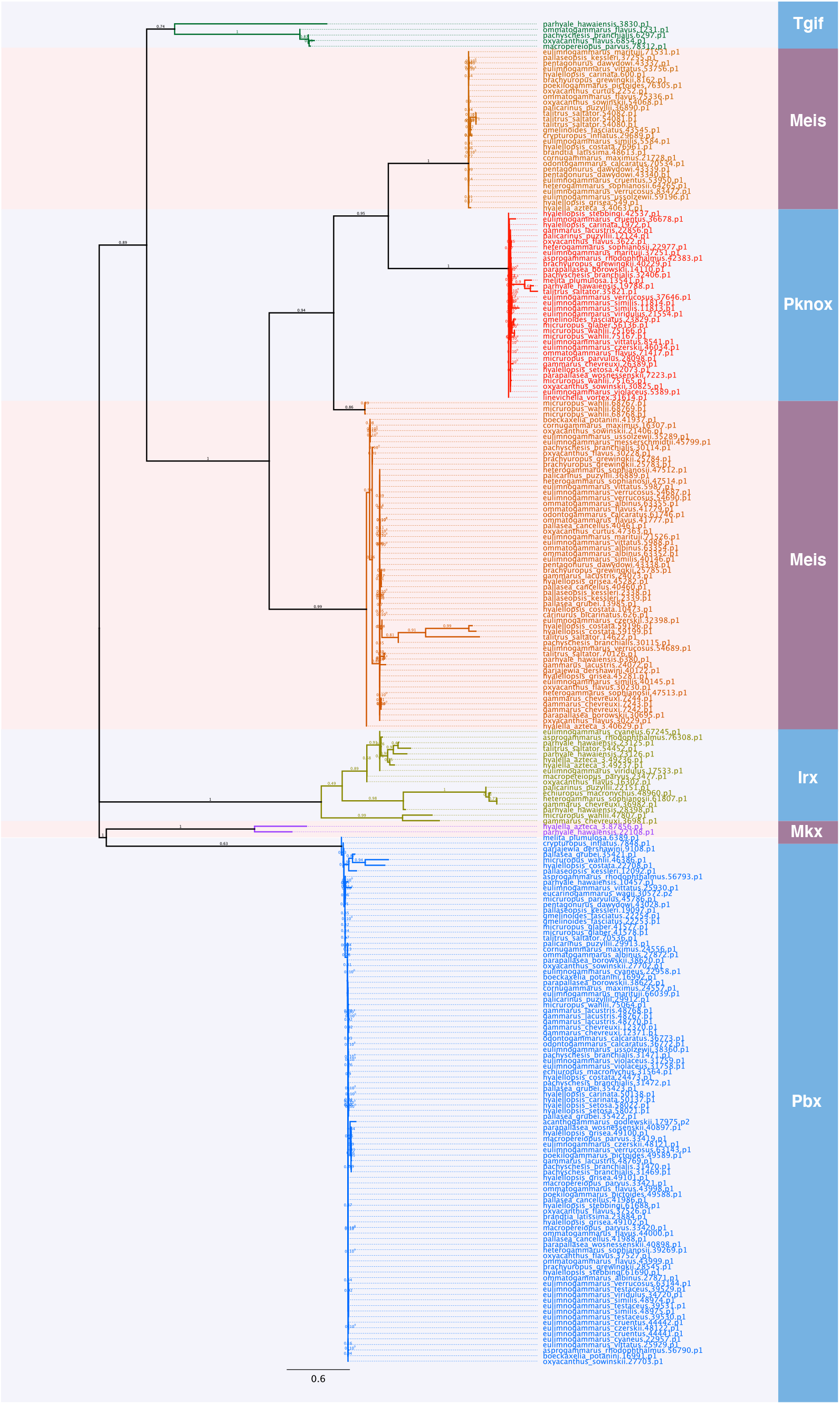
Maximum likelihood tree of TALE class genes in amphipod crustaceans.

## Conclusion

TALE class genes identified from amphipod crustaceans demonstrated conserved patterns of evolution. Availability of the gene sequences may contribute to current and future research focusing on elucidating and understanding evolutionary processes in non-model organisms.

## Notes

### Competing Interest Statement

The authors have declared no competing interest.

